# *The Crown Pearl*: a draft genome assembly of the European freshwater pearl mussel *Margaritifera margaritifera* (Linnaeus, 1758)

**DOI:** 10.1101/2020.12.06.413450

**Authors:** André Gomes-dos-Santos, Manuel Lopes-Lima, André M. Machado, António Marcos Ramos, Ana Usié, Ivan N. Bolotov, Ilya V. Vikhrev, Sophie Breton, L. Filipe C. Castro, Rute R. da Fonseca, Juergen Geist, Martin E. Österling, Vincent Prié, Amílcar Teixeira, Han Ming Gan, Oleg Simakov, Elsa Froufe

## Abstract

Since historical times, the inherent human fascination with pearls turned the freshwater pearl mussel *Margaritifera margaritifera* (Linnaeus, 1758) into a highly valuable cultural and economic resource. Although pearl harvesting in *M. margaritifera* is nowadays residual, other human threats have aggravated the species conservation status, especially in Europe. This mussel presents a myriad of rare biological features, e.g. high longevity coupled with low senescence and Doubly Uniparental Inheritance of mitochondrial DNA, for which the underlying molecular mechanisms are poorly known. Here, the first draft genome assembly of *M. margaritifera* was produced using a combination of Illumina Paired-end and Mate-pair approaches. The genome assembly was 2,4 Gb long, possessing 105,185 scaffolds and a scaffold N50 length of 288,726 bp. The *ab initio* gene prediction allowed the identification of 35,119 protein-coding genes. This genome represents an essential resource for studying this species’ unique biological and evolutionary features and ultimately will help to develop new tools to promote its conservation.

## 1. Introduction

Pearls are fascinating organic gemstones that have populated the human beauty imaginary for millennia. Legend says that the famous Egyptian ruler Cleopatra, to display her wealth to her lover Marc Antony, dissolved a pearl in a glass of vinegar and drank it. The human use of pearls or their shell precursor material, the mother-of-pearl (nacre), is ancient. The earliest known use of decorative mother-of-pearl dates to 4200 BC in Egypt, with pearls themselves only becoming popular around 600 BC. Before the arrival of marine pearls to Europe, most were harvested from a common and widespread freshwater bivalve, the freshwater pearl mussel *Margaritifera margaritifera* L. 1758, where generally one pearl is found per 3,000 mussels leading to massive mortality^1^. In Europe, during the Roman Empire period, pearls were a desirable luxury, so that it is believed that one of the reasons that persuaded Julius Caesar to invade Britain was to access its vast freshwater pearl resources^2^. *Margaritifera margaritifera* freshwater pearls were extremely valuable being included in many royal family jewels such as the British, Scottish, Swedish, Austrian and German crown jewels and even in the Russian city’s coat of arms^2–5^. Although over-harvesting represented a serious threat to the species for centuries (mostly in Europe and Russia), there has been a decrease in interest and demand for freshwater pearls in the 20^th^ century^4^. However, the global industrialization process introduced stronger threats to the survival of the species^6–8^. In fact, *M. margaritifera* belongs to one of the most threatened taxonomic groups on earth, the Margaritiferidae^6^. The species was once abundant in cool oligotrophic waters throughout most of northwest Europe and northeast North America^6–8^. However, habitat degradation, fragmentation and pollution have resulted in massive population declines^8^. Consequently, the Red List of Threatened Species from the International Union for Conservation of Nature (IUCN) has classified *M. margaritifera* as Endangered globally and Critically Endangered in Europe^7,9^. Population declines are particularly concerning in Europe, where a lot of investment has been done in rehabilitation and propagation projects aimed at improving the species conservation status^9,10^. North America and Russia seem to be able to control populations sizes by maintaining more isolated and less threat exposed populations^7^.

Besides being able to produce pearls, *M. margaritifera* presents many other remarkable biological characteristics, e.g. is among the most longest-living invertebrates, reaching up to 280 years^6,11^; displays very weak signs of senescence, referred as the concept of “*negligible senescence*”^12^; has an obligatory parasitic larval stage on salmonid fishes used for nurturing and dispersion^8,10^; and, like many other bivalves (see Gusman et al.^13^ for a recent enumeration), shows an unusual mitochondrial DNA inheritance system, called Doubly Uniparental Inheritance or DUI^14,15^. Although these biological features are well described, the molecular mechanisms underlying their regulation and functioning are poorly studied and practically unknown.

Thus, a complete genome assembly for *M. margaritifera* is critical for developing the molecular resources required to improve our knowledge of such mechanisms. These resources can then be used in multiple fields, such as in conservation biology (e.g. to overcome the main bottlenecks of propagation programs); in freshwater pearl production industry (e.g. to better understand biomineralization mechanisms); in biomedical applications (e.g. to study bone regeneration); and in ageing and senescence studies among others.

In the last decade, the decreasing cost of next generation sequencing, coupled with improved bioinformatic tools, has facilitated the generation of genomic resources for non-model organisms. Several Mollusca genomes are currently available and new assemblies are released every year at an increasing trend (reviewed in^16–18^). Despite this, to date, only two Unionida mussel genomes have been published, those of *Venustaconcha ellipsiformis* (Conrad, 1836)^19^ and *Megalonaias nervosa* (Rafinesque, 1820)^20^. These represent valuable comparative resources and are among the largest bivalve genomes sequenced to date (1.80 Gb and 2.36 Gb respectively)^16,18^; known Bivalvia genome sizes range from 0.559 Gb (*Crassostrea gigas*, Ostreida^21^) to 2.38 Gb *(Modiolus philippinarum*, Mytiloida^22^). Unlike many other bivalves (e.g.^23,24^), both *V. ellipsiformis* and *M. nervosa* genomes revealed relatively low levels of heterozygosity, with estimated values per site of 0.0060 and 0.0077, respectively^19,20^.

The current study presents the first draft genome assembly of the freshwater pearl mussel *M. margaritifera*. The genome assembly was performed combining Illumina Paired-End short reads with Illumina Mate-Paired reads. The assembled genome has a total length of 2,4 Gb distributed throughout 105,185 scaffolds, with a GC content of 35.42% and scaffold N50 length of 288,726 bp. More than half of the genome was found to be composed of repetitive elements (i.e. 59.07%) and 35,119 protein coding genes were predicted in this initial annotation.

## 2. Material and Methods

### 2.1. Sample collection, DNA extraction and sequencing

One *M. margaritifera* (Linnaeus, 1758) specimen was collected from the River Tua, Douro basin in the North of Portugal (permit 284/2020/CAPT and fishing permit 26/20 issued by ICNF - Instituto de Conservação da Natureza e das Florestas). The whole individual is stored in 96% ethanol at the Unionoid DNA and Tissue Databank, CIIMAR, University of Porto. Genomic DNA (gDNA) was extracted from the foot tissue using DNeasy Blood and Tissue Kit (Qiagen, Hilden, Germany) according to the manufacturer’s instructions.

Two distinct NGS libraries and sequencing approaches were implemented i.e. Illumina Paired-end reads (PE) and Illumina long insert size Mate-pair reads (MP). For Illumina PE library preparation, approximately 100ng of gDNA as measured using Qubit Broad-Range Kit (Invitrogen, Santa Clara, CA, USA) was sheared to approximately 300-400 bp using the Qsonica Q800R system (Qsonica, Newton, CT, USA). The sheared DNA was then prepared for sequencing using NEB Ultra DNA kit with standard Illumina adapter and sequenced in an Illumina machine NovaSEQ6000 system located at Deakin Genomics Centre using a run configuration of 2×150 bp. Illumina MP library preparation and sequencing were performed by Macrogen Inc., Korea, where a 10kb insert size Nextera Mate Pair Library was constructed and subsequently sequenced in a NovaSeq6000 S4 using a run configuration of 2×150 bp.

### 2.2. Genome size and heterozygosity estimation

The overall characteristics of the genome were accessed using PE reads. Firstly, the general quality of the reads was evaluated with FastQC (https://www.bioinformatics.babraham.ac.uk/projects/fastqc/). The raw reads were then quality trimmed with Trim Galore v.0.4.0 (https://www.bioinformatics.babraham.ac.uk/projects/trim_galore/), allowing the trimming of adapter sequences and removal of low-quality reads using Cutadapt^25^ followed by another quality check with FastQC. Clean reads were used for genome size estimation in a two-step approach: (i) using Jellyfish v.2.2.10 for counting and histogram construction of k-mer frequency distributions, with k-mers length of 25 and 31 and (ii) using histograms of frequency distribution to estimate the genome size, heterozygosity rate and repeat content of the genome, with GenomeScope2^26,27^.

### 2.3. Genome assembly and quality assessment

Long range Illumina mate paired reads quality processing was as described above and both PE and MP cleaned reads were used for whole genome assembly. The assembly was produced by running Meraculous v.2.2.6 with several distinct k-mer sizes (meraculoususing)^28^. This allowed determining the optimal kmer size of 101. Genome assembly metrics were estimated using QUAST v5.0.2^29^. Assembly completeness, heterozygosity and collapsing of repetitive regions were evaluated through analysis of k-mer distribution using PE reads, with the tool “KAT comp” from the K-mer analysis toolkit^30^ Furthermore, PE reads were aligned to the genome assembly using BBMap^31^. BUSCO v. 3.0.2^32^ was used to provide a quantitative measure of the assembly completeness, with a curated list (i.e. OrthoDB) of near-universal single-copy orthologs. Here, both eukaryotic (303 single-copy orthologs) and metazoan (978 single-copy orthologs) libraries profiles were used to test the genome assembly completeness.

### 2.4. Repeat Sequences and Gene Models predictions

Given the generally high composition of repetitive elements in Mollusca genomes (e.g.^16^) they should be identified and masked before proceeding to genome annotation. An annotated library of repetitive elements was created for *M. margaritifera* genome assembly, using RepeatModeler v.2.0.1.^33^ (excluding sequences <2.5 kb). Afterwards, repetitive elements were soft masked using RepeatMasker v.4.0.7.^34^ combining the repetitive elements for the taxa “Bivalvia”, from the RepeatMasker database (comprising the databases Dfam_consensus-20170127 and RepBase-20181026), with the newly constructed library for *M. margaritifera* genome.

BRAKER2 pipeline v2.1.5^35,36^ was used for gene prediction in the genome. First, all RNA-seq data of *M. margaritifera*^37,38^ available on GenBank were downloaded, assessed with FastQC v.0.11.8 (http://www.bioinformatics.babraham.ac.uk/projects/fastqc/), quality-controlled and uniformized with Trimmomatic v.0.38^39^ (Parameters, LEADING:5 TRAILING:5 SLIDINGWINDOW:4:20 MINLEN:36), the sequencing errors corrected with Rcorrector v.1.0.3. Afterwards, the RNA-seq data was aligned to the masked genome assembly, using Hisat2 v.2.2.0 with the default settings^40^. Secondly, the complete proteomes of 13 mollusc species, one Chordata (*Ciona intestinalis*) and one Echinodermata (*Strongylocentrotus purpuratus*) were downloaded from distinct public databases (Supplementary Table S1) and used as additional evidence for gene prediction. The BRAKER2 pipeline was applied with the parameters (--etpmode; --softmasking; --UTR=off; --crf; --cores=30) and following the authors’ instructions^35,36^. The resulting gene predictions (i.e. gff3 file) were renamed, cleaned and filtered using AGAT v.0.4.0^41^, correcting coordinates of overlapping gene prediction, removing predicted coding sequence regions (CDS) with less than 100 amino acid (in order to avoid a high rate of false positive predictions) and removing incomplete gene predictions (i.e. without start and/or stop codons). Functional annotation was first conducted by searching for protein domain information using InterProScan v.5.44.80^42^, and afterwards, a protein blast search was conducted using DIAMOND v. 0.9.32^43^ against SwissProt (Download at 2/07/2020), TREMBL (Download at 2/07/2020) and RefSeq-NCBI (Download at 3/07/2020)^44,45^.

### 2.5. Phylogenetic analyses

For the phylogenetic assessment, the proteomes of 12 molluscan species were downloaded from distinct public databases (Supplementary Table S2) and included 11 Autobranchia bivalves and two outgroup species, i.e. the Cephalopoda *Octopus bimaculoides* and Gastropoda *Biomphalaria glabrata* (Figure 3). To retrieve single-copy orthologs between these 12 species and *M. margaritifera*, the protein sets were first clustered into families, using OrthoFinder v2.4.0^46^ specifying msa as the method of gene tree inference (-M). The resulting 118 single copy orthologous sequences were individually aligned using MUSCLE v3.8.31^47^, with default parameters and subsequently trimmed with TrimAl v.1.2^48^ specifying a gap threshold of 0.5 (gt). Trimmed sequences were then concatenated using FASconCAT-G (https://github.com/PatrickKueck/FASconCAT-G). The best molecular evolutionary model was estimated using ProTest v.3.4.1^49^. Phylogenetic inferences were conducted in IQ-Tree v.1.6.12^50^ for Maximum Likelihood analyses (with initial tree searches followed by ten independent runs and 10000 ultra-bootstrap replicates) and MrBayes v.3.2.6^51^ for Bayesian Inference (two independent runs, 1,000,000 generations, sampling frequency of one tree per 1000 generations). All phylogenetic analyses were applied using the substitution model LG+I+G.

### 2.6. Hox and ParaHox gene identification and phylogeny

To identify the repertoire Hox and ParaHox genes *in M. margaritifera*, a similarity search by BLASTn^52^ of the CDS of *M. margaritifera* genome, was conducted using the annotated homeobox gene set of *Crassostrea gigas*^53,54^. Candidate CDSs were further validated for the presence of the homeodomain by CD-Search^55^. Finally, each putative CDS identity was verified by BLASTx and BLASTp^52^ searches in Nr-NCBI nr database and phylogenetic analyses. Since the search was conducted in the annotated genome (i.e. scaffolds over 2.5kb), when genes were not found, a new search was conducted in the remaining scaffolds. At the end, any genes still undetected were search in the Transcriptome assembly of the species (Bioproject: PRJNA369722)^37^. Due to the phylogenetic proximity and for comparative purposes, Hox and ParaHox genes were also searched in the genome assembly of *Megalonaias nervosa*^20^.

For phylogenetic assessment of Hox and Parahox genes, amino acid sequences of homeodomain of the genes from *M. margaritifera* and *M. nervosa*, were aligned with other Mollusca orthologs (^56,57^ and references within; Supplementary File1). Molecular evolutionary models and Maximum Likelihood phylogenetic analyses were obtained using IQ-TREE v.1.6.12^50,58^.

## 3. Results and Discussion

### 3.1. Sequencing Results

A total of 494 Gb (~209x) of raw PE and 76 Gb (~32x) of raw MP data were generated, which after trimming and quality filtering were reduced by 0.3% and 10% respectively (Table 1). GenomeScope2 model fitting of the k-mer distribution analysis estimated a genome size between 2.31-2.36 Gb and very low heterozygosity between 0.127-0.105% (Figure 2). Although larger than the genome of *V. ellipsiformis* (i.e. 1.80 Gb), the size estimation of the *M. margaritifera* genome is in line with the recently assembled Unionida mussel *M. nervosa*^20^ (i.e. 2.38 Gb). The estimated heterozygosity is the lowest observed within Unionida genomes^19,20^ and one of the lowest in Mollusca^16^, which is remarkable considering it refers to a wild individual. This low value is likely a consequence of population bottlenecks during glaciations events, which have been shown to shape the evolutionary history of many freshwater mussels (e.g.^19,59,60^) and may also be enhanced by recent human-mediated threats.

**Figure 1.**
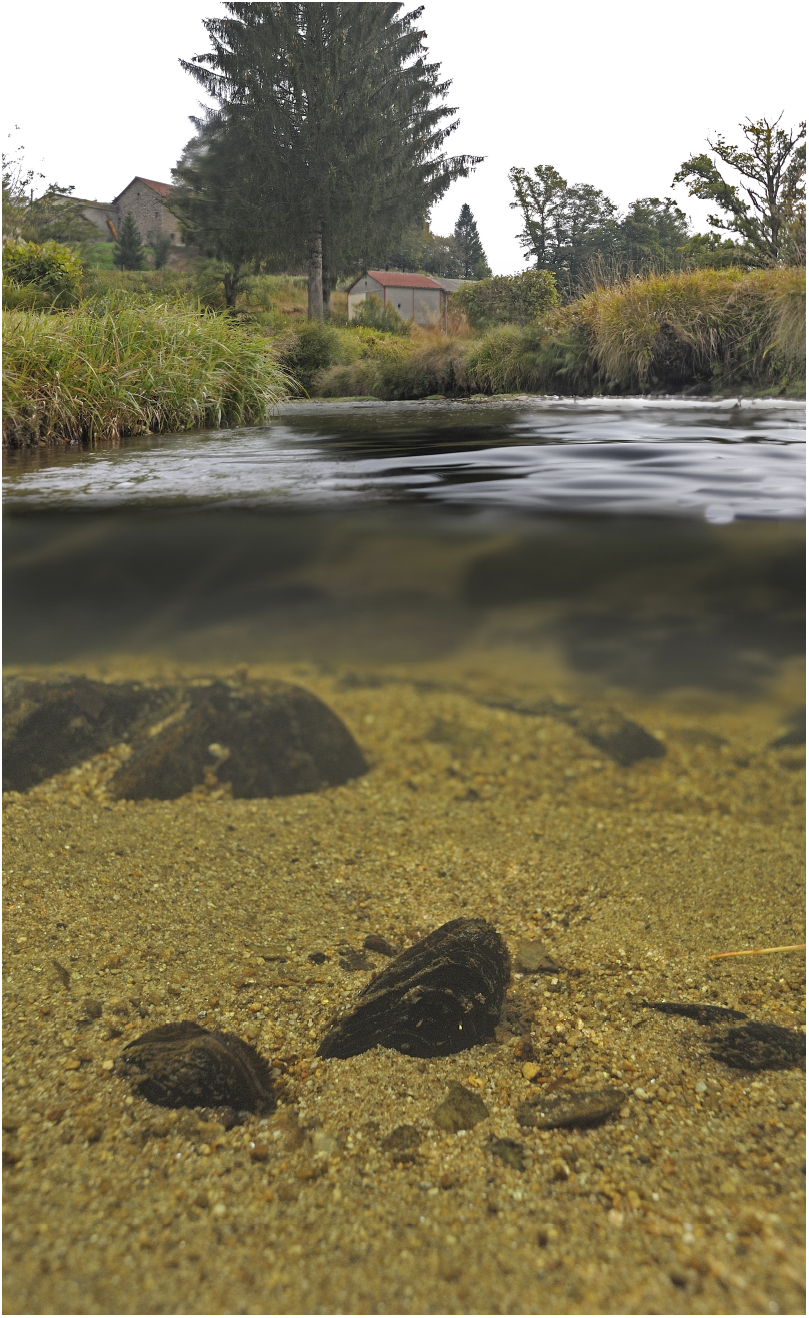
*Margaritifera margaritifera* specimen in its natural habitat.

**Figure 2.**
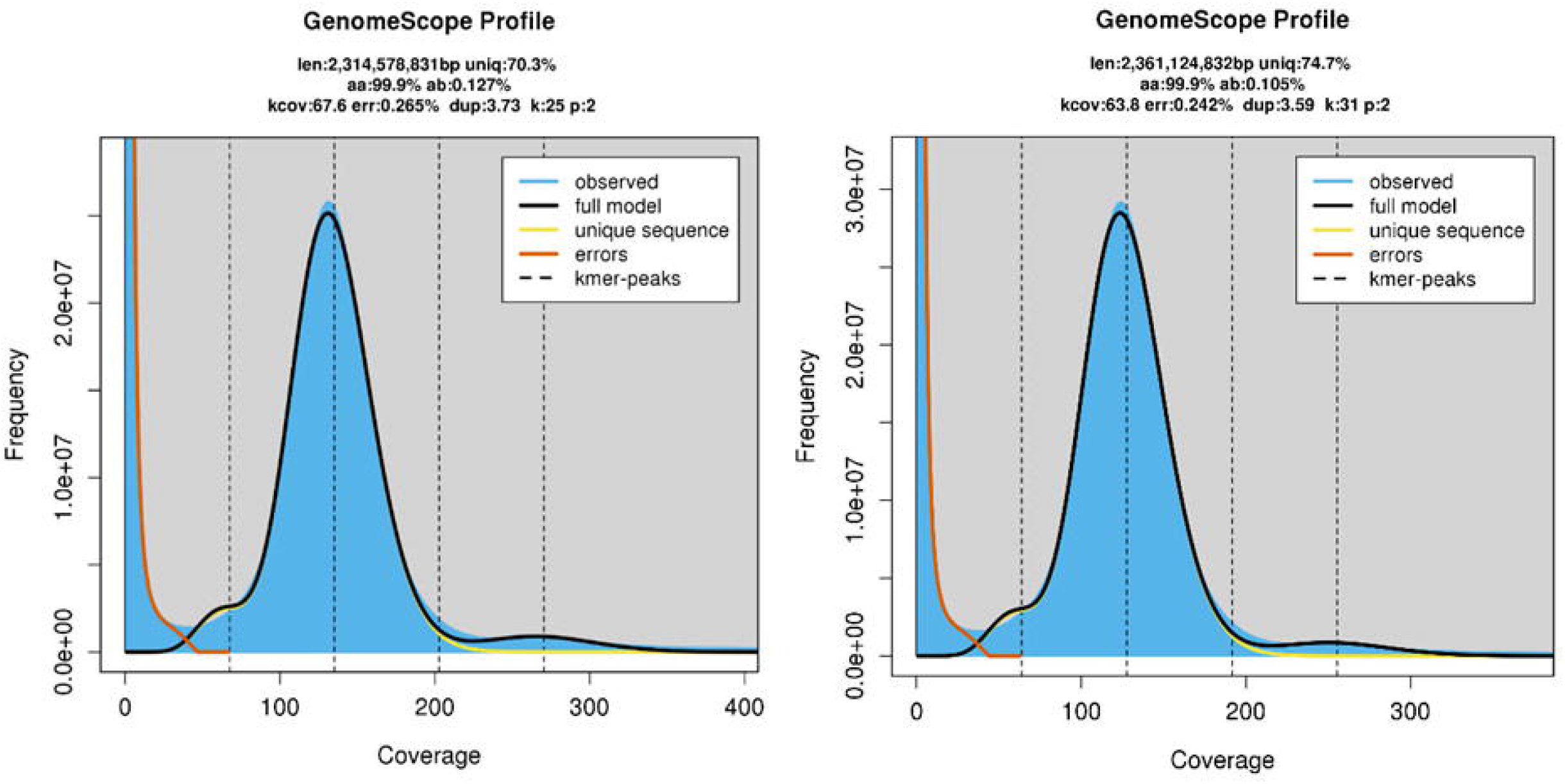
GenomeScope2 k-mer (25 and 31) distribution displaying estimation of genome size (len), homozygosity (aa), heterozygosity (ab), mean kmer coverage for heterozygous bases (kcov), read error rate (err), the average rate of read duplications (dup), k-mer size used on the run (k:) and ploidy (p:).

**Table 1.**
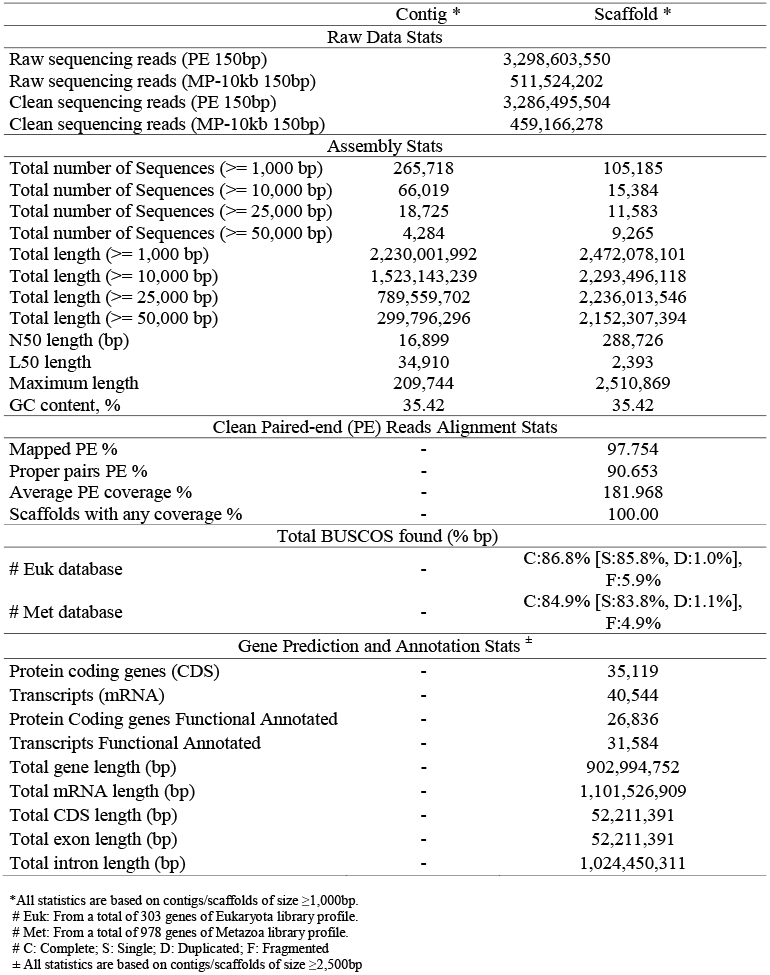
*Margaritifera margaritifera* sequencing, genome assembly, read alignment, gene prediction and annotation general statistics.

### 3.2. *Margaritifera margaritifera* de novo genome assembly

The Meraculous assembly and scaffolding yield a final genome size of 2.47 Gb with a contig N50 of 16,899 bp and a scaffold N50 of 288,726 bp (Table 1). Both N50 values are significantly higher than *V. ellipsiformis* genome assembly, i.e. 3,117 bp and 6,523 bp, respectively^19^. Presently, this *M. margaritifera* genome assembly reveals the highest scaffold N50 of the three Unionida genomes currently available^19,20^. On the other hand, *M. nervosa* genome assembly contig N50, i.e. 51,552 bp, is higher than *M. margaritifera*, which is expected given the use of Oxford Nanopore ultra-long reads libraries in the assembly produced by Rogers et al^20^. BUSCOs scores of the final assembly indicate a fairly complete genome assembly (Table 1) and although the contiguity is lower when compared with other recent Bivalve genome assemblies, the low percentage of fragmented genes (i.e. 5.9% for Eukaryota and 4.9% for Metazoa) gives further support to the quality of the genome assembly. Similarly, the slight difference observed between the genome size and the initial size estimation is unlikely to be a consequence of erroneous assembly duplication, as duplicated BUSCOs scores are also low (i.e. 1% for Eukaryota and 1.1% for Metazoa). The quality of the genome assembly is further supported by the high percentages of PE reads mapping back to the genome (i.e. 97.75%, Table 1), as well as the KAT k-mer distribution spectrum (Figure 3), which demonstrates that almost no read information was excluded from the final assembly. Overall, these statistics indicate that the *M. margaritifera* draft genome assembly here presented is fairly complete, nonredundant, and useful resource for various applications.

**Figure 3.**
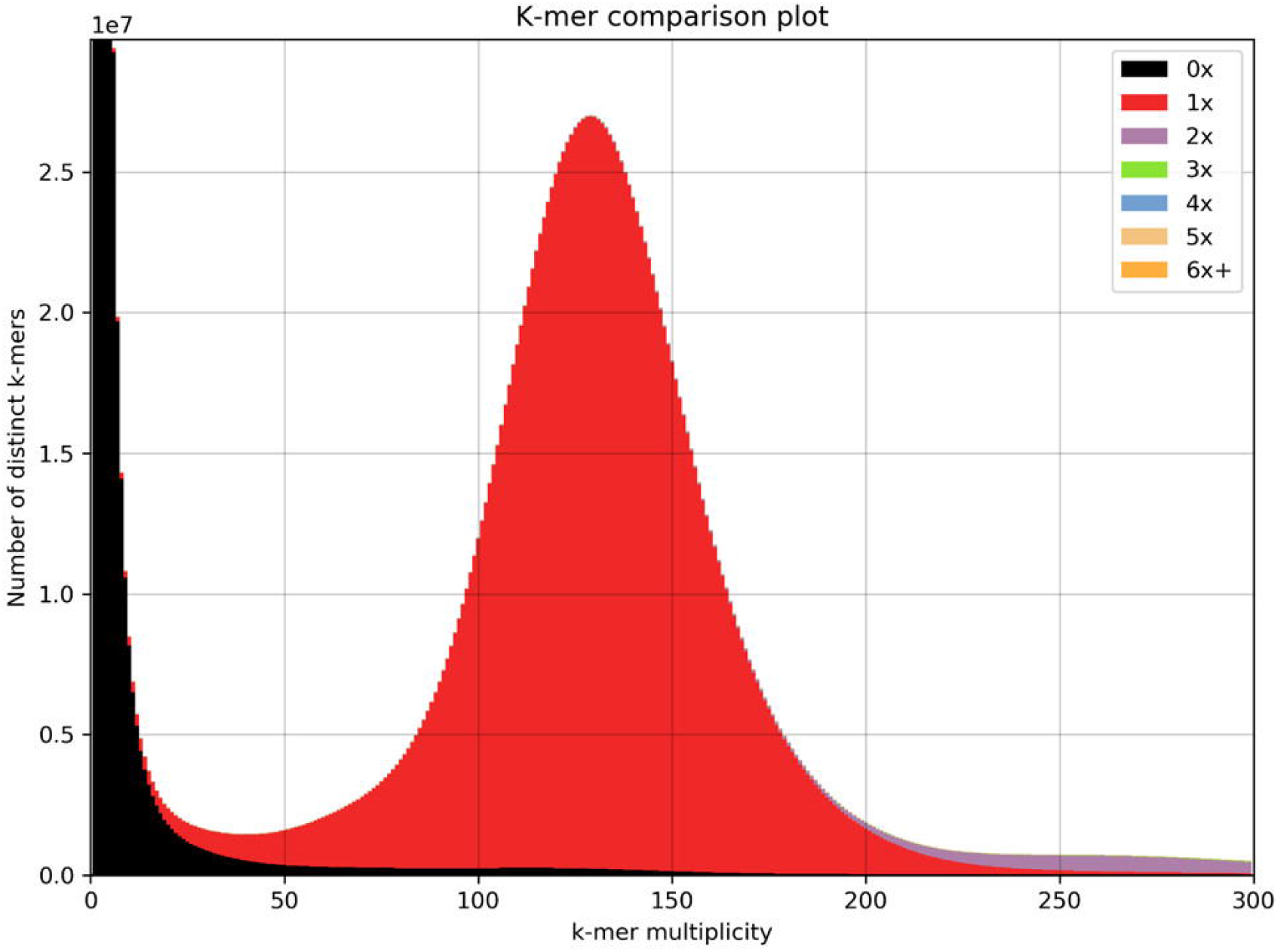
*Margaritifera margaritifera* genome assembly assessment using KAT comp tool to compare the Illumina Paired-end reads k-mer content within the genome assembly. Different colours represent the read k-mer frequency in the assembly.

### 3.3. Repeat Identification and Masking and Gene Models Prediction

The use of the custom repetitive library combined with the RepBase^61^ “Bivalvia” library, resulted in masking repetitive elements in more than half of the genome assembly, i.e. 59.07% (Table 2). Most of the annotated repetitive elements were unclassified (31.86%), followed by DNA elements (16.00%), long interspersed nuclear elements (LINEs) (6.13%), long terminal repeats (LTRs) (3.72%) and short interspersed nuclear elements (SINEs) (0.79%). After masking, gene prediction resulted in the identification of 35,119 protein-coding genes, with an average gene length of 25,712 bp and average CDS length of 1,287 bp (Supplementary Table S3). Furthermore, 26,836 genes were functionally annotated by similarity to at least one of the three databases used in the annotation (Table 1). The number of predicted genes is in accordance to those observed in other bivalves (and Mollusca) genome assemblies, which although highly variable, in average have around 34,949 predicted genes (calculated from Table 2 of Gomes-dos-Santos et al.^16^). Although the number of genes predicted within the three Unionida genomes is highly variable, i.e. 123,457 in *V. ellipsiformis*, 49,149 in *M. nervosa* and 35,119 in *M. margaritifera*, a direct comparison should be taken with caution, given the considerable differences in genome qualities and the different gene predictions strategies applied in the three assemblies.

**Table 2.**
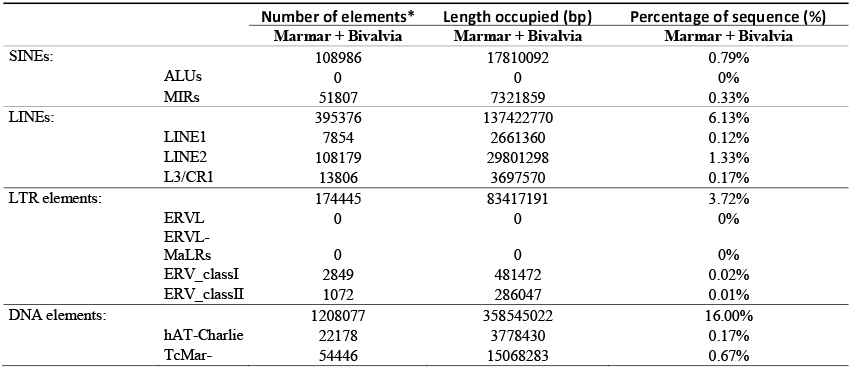
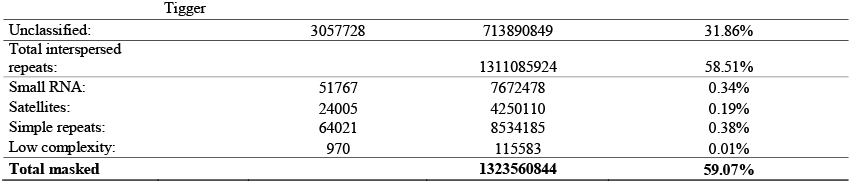
Statistics of the content of repetitive elements in the *M. margaritifera* genome assembly. Values were produced by RepeatMasker using a RepeatModeler’s custom build *M. margaritifera* repeat library (abbreviated with “Marmar”) combined with the RepBase Biavalve repeat library (RepeatMasker option-lib).

### 3.4. Single Copy Orthologous Phylogeny

Both Maximum Likelihood and Bayesian Inference phylogenetic trees revealed the same topology with high support for all nodes (Figure 3). The phylogeny recovered the reciprocal monophyletic groups Pteriomorphia (represented by Orders Ostreida, Mytilida, Pectinida and Arcida) and Heteroconchia (represented by Orders Unionida and Venerida). These results are in accordance with recent comprehensive bivalve phylogenetic studies^38,62–64^. The only difference is observed within Pteriomorphia, where two sister clades are present, one composed by Arcida and Pectinida and the other by Mytilida and Osteida (Figure 3), while accordingly to the most recent phylogenomic studies, Arcida appears basal to all other Pteriomorphia^38,63,64^. It is noteworthy that Arcida and Pectinida clade is the less supported in the phylogeny, which together with the fact that many Pteriomorphia clades are missing in the present study, should explain these discrepant results. Heteroconchia is divided into monophyletic Palaeoheterodonta and Heterodonta (here only represented by two Euheterodonta bivalves). As expected, the two Unionida species, i.e. *M. nervosa* and the newly obtained *M. margaritifera*, are placed within Palaeoheterodonta.

### 3.5. Hox and ParaHox gene repertoire and phylogeny

Homeobox genes refer to a family of homeodomain-containing transcription factors with important roles in Metazoan development by specifying anterior-posterior axis and segment identity (e.g.^65,66^). Many of these genes are generally found in tight evolutionary conserved physical clusters (e.g.^67,68^). Hox genes are typically arranged into tight physical clusters, showing temporal and spatial collinearity^69^. Consequently, Hox genes provide useful information for understanding the emergence of morphological novelties, understanding the historical evolution of the species, infer ancestral genomic states of genes/clusters and even study genome rearrangements, such as whole-genome duplications (e.g.^65,66,70^). Given the disparate body plans in molluscan classes, the study of Hox cluster composition, organization and gene expression has practically become a standard in Mollusca genome assembly studies^21,22,78–82,57,71–77^. Homeobox genes are divided into four classes, of which the Antennapedia (ANTP)-class (Hox, ParaHox, NK, Mega-homeobox, SuperHox) is the best studied, particularly the Hox and ParaHox clusters^57,70,75^. The number of genes from these two clusters is relatively well conserved across Lophotrochozoa, with Hox cluster being composed of 11 genes (3 anterior, 6 central and 2 posterior) and ParaHox cluster composed of 3 genes. Although several structural and compositional differences have been observed within Mollusca ANTP-class (e.g. Bivalvia^21^, Cephalopoda^72^, Gastropoda^74^ and Polyplacophora^80^, most Bivalvia seem to retain the gene composition expected for lophotrochozoans: Hox1, Hox2, Hox3, Hox5, Lox, Antp, Lox4, Lox2, Post2, Post1 for the Hox cluster and Gsx, Xlox, Cdx for the ParaHox cluster^81^. Consequently, the identification of these genes on a bivalve genome assembly represent further validation of the genome completeness and overall correctness. Furthermore, to the best of our knowledge, this study reports for the first time the Hox and ParaHox genes were identified Unionida. A single copy of the 3 ParaHox and 10 Hox genes were found in the *M. margaritifera* genome assembly (Supplementary Table S4). Despite an intensive search, no evidence of the presence of Hox4 was detected. However, the gene was identified in the *M. margaritifera* transcriptome, thus confirming its presence in the species. All genes, apart from *Antp* and Lox5, were scattered in different scaffolds, with Hox5, Post1 and Gsx being present in scaffolds smaller than 2.5kb (Supplementary Table S4). Both the small proximity between Antp and Lox5 and the fact that both genes are expressed in the same direction are in accordance with the results observed in other bivalves, including in the phylogenetically closest species (from which Hox cluster has been characterized), i.e. the Venerida clam *Cyclina sinensis* (Gmelin, 1791)^57^. The fact that the remaining genes were scattered in the different scaffolds is likely a consequence of the low contiguity of the genome assembly since the distances between Bivalvia Hox genes within a cluster can be as high as 9.9 Mb^57^. Conversely, 3 Hox and 1 ParaHox genes were found in the *M. margaritifera* transcriptome assembly and 9 Hox and 1 ParaHox gene were found in *M. nervosa* genome assembly (Supplementary Table S4). Finally, to further validate the identity of the identified Hox and ParaHox genes, a phylogenetic analysis using the homeodomains (encoded 60–63 amino acid domain) of several Mollusca species was conducted (Figure 5). All Hox and ParaHox genes of *M. margaritifera* (as well as *M. nervosa*) were well positioned within their respective orthologous genes from other Mollusca species (Figure 4), thus confirming their identity.

**Figure 4.**
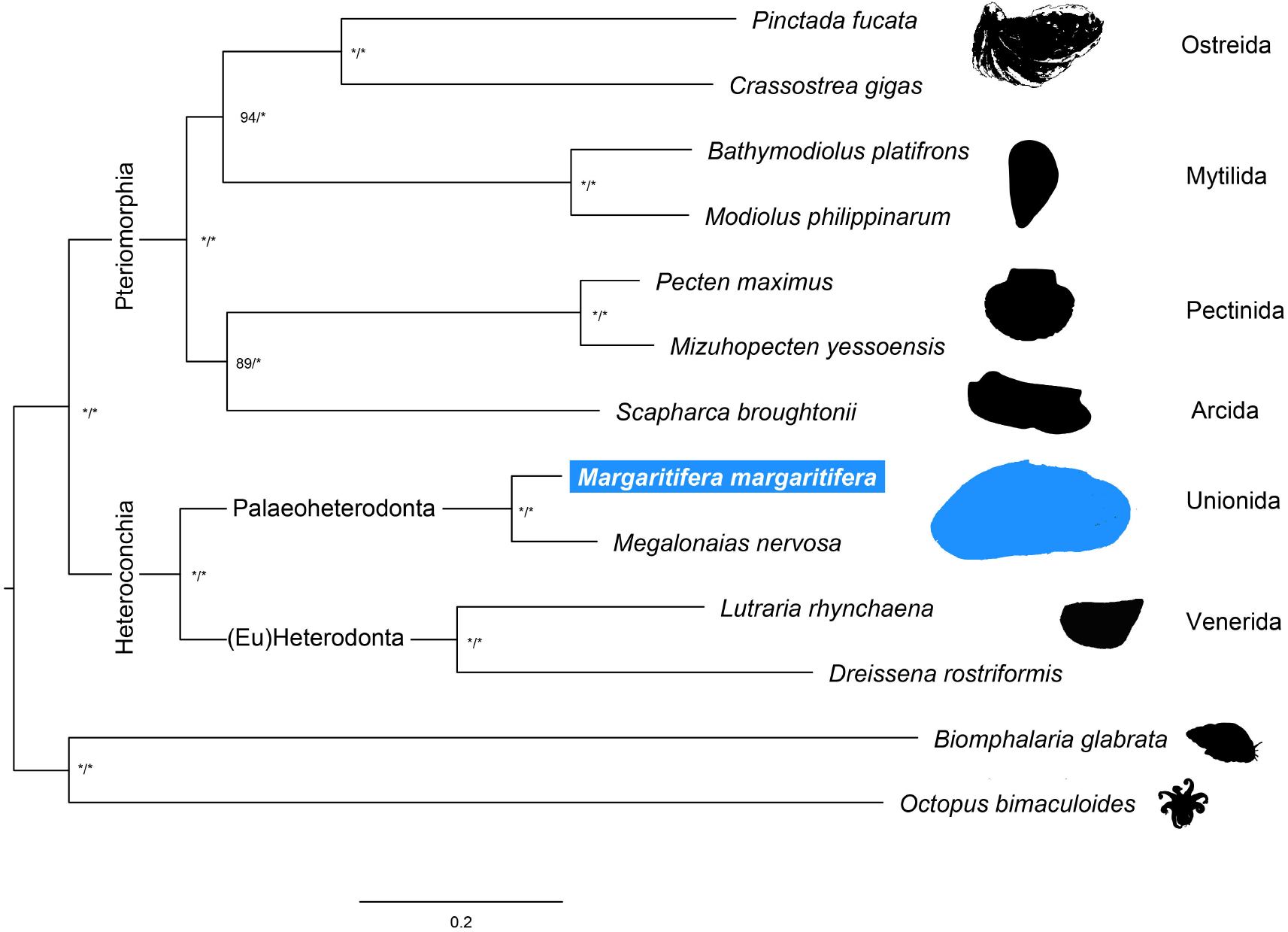
Maximum Likelihood phylogenetic tree based on concatenated alignments of 118 single copy orthologous amino acid sequences retrieved by OrthoFinder. * above the nodes refer to bootstrap and posterior probabilities support values above 99%.

**Figure 5.**
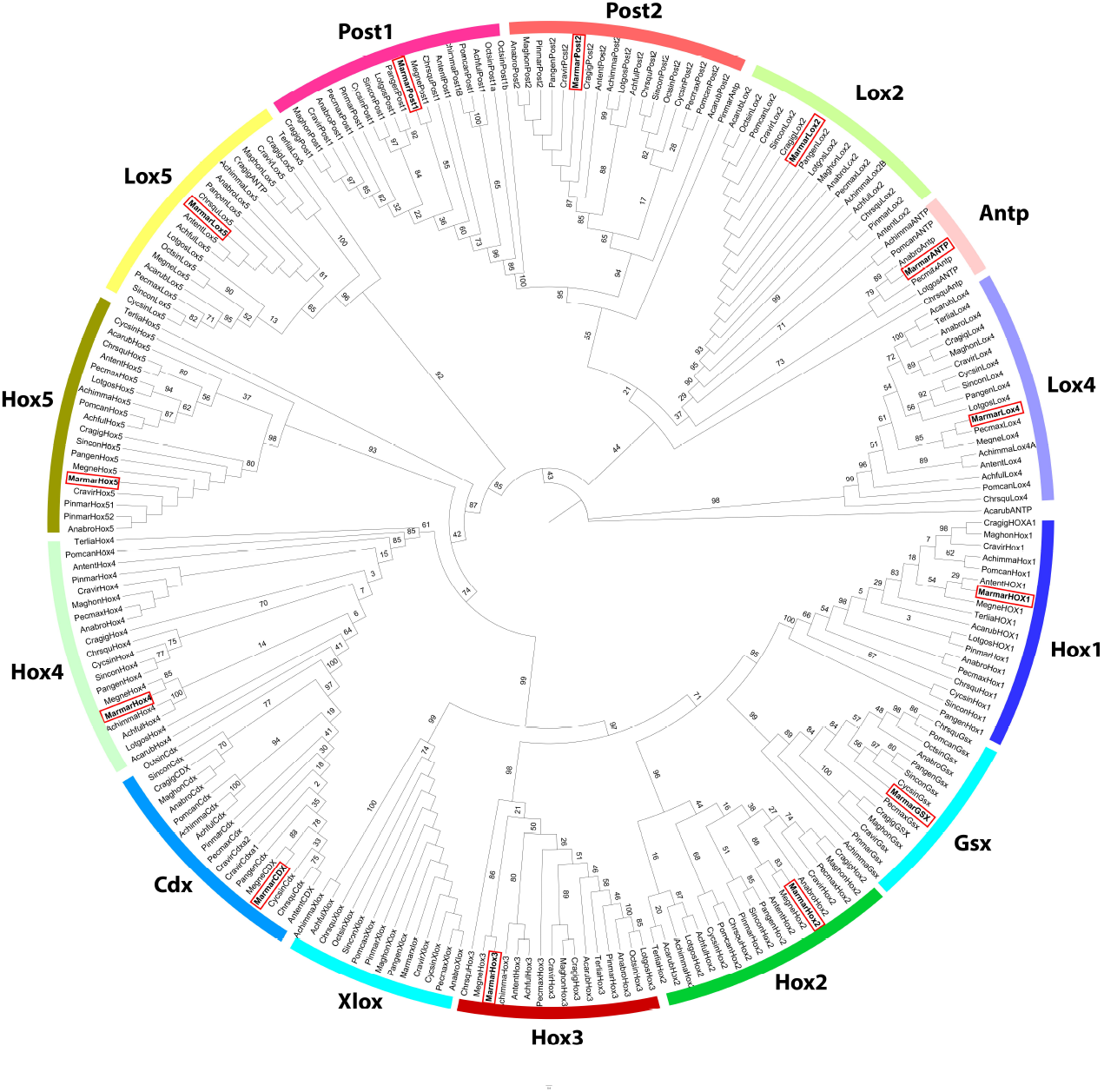
Hox and ParaHox Maximum Likelihood gene tree constructed using Mollusca homeodomain amino acid sequences. Bootstrap values are presented above the nodes.

### 3.6. Conclusion and future perspectives

Unionida freshwater mussels are a worldwide distributed and diverse group of organisms with 6 recognized families and around 800 described species^83,84^. These organisms play fundamental roles in ecosystems, such as water filtration, nutrient cycling and sediment bioturbation and oxygenation^85,86^, allowing to maintain and support freshwater communities^10^. However, as a consequence of several anthropogenic threats, freshwater mussels are experiencing a global-scale decline^10,87^. *Margaritifera margaritifera* belongs to the most threatened of the 6 Unionida families, i.e. Margaritiferidae. Despite all this, our understanding of the genetics of this species is still to date restricted to a few mtDNA markers phylogenetic and restricted phylogeographical studies^6,88–90^ as well as neutral genetic markers (SSR)^89,91,92^, making the availability of the present genome a timely resource. Being the first representative genome of the family Margaritiferidae, it will help launch both basic and applied genomic-level research on the unique biological and evolutionary features characteristic of this emblematic group.

## Funding

AGS was funded by the Portuguese Foundation for Science and Technology (FCT) under the grant SFRH/BD/137935/2018. This research was developed under ConBiomics: the missing approach for the Conservation of freshwater Bivalves Project No NORTE-01-0145-FEDER-030286, co-financed by COMPETE 2020, Portugal 2020 and the European Union through the ERDF, and by FCT through national funds. Additional strategic funding was provided by FCT UIDB/04423/2020 and UIDP/04423/2020. Authors interaction and writing of the paper was promoted and facilitated by the COST Action CA18239: CONFREMU - Conservation of freshwater mussels: a pan-European approach.

## Data Availability

All the raw sequencing data are available from GenBank via the accession numbers SRR13091478, SRR13091479 and SRR13091477. The assembled genomes are available in the assession number JADWMO000000000, under the BioProject PRJNA678877 and BioSample SAMN16815977 (Supplementary Table S5). Fasta alignment of homeodomain amino acid sequences from Hox and ParaHox genes used in gene tree construction is available in Additional File 1. The scaffolds in which homeodomains were detected (as described in Supplementary Table S4) are available as Supplementary File 2. The repeat masked genome assembly, BRAKER2 prediction statistic and prediction gff files, as well as all predicted genes, transcripts and amino acid sequence files are available at Figshare: https://doi.org/10.6084/m9.figshare.13333841

## Conflict of interest

None declared.

## Supplementary data

Supplementary Table S1 – List of proteomes used for BRAKER2 gene prediction pipeline.

Supplementary Table S2 – List of proteomes used to retrieve single-copy orthologs in OrthoFinder v2.4.0.

Supplementary Table S3 – BRAKER2 gene prediction complete report.

Supplementary Table S4 – Genomic locations of Hox and ParaHox genes in the genome assemblies of *M. margaritifera* and *M. nervosa* and trancriptome assembly of *M. margaritifera*.

Supplementary Table S5 – Descriptors and acession numbers of tissue samples, raw data and assemblies of *Margaritifera margaritifera*

Supplementary File 1 – Fasta alignment of homeodomain amino acid sequences from Hox and ParaHox genes used in gene tree construction. Sequences used include the Hox and ParaHox homeodomains obtained in the current study as well as other Mollusca homeodomain sequences retrieved from (Huan et al., 2020; Li et al., 2020) and references within.

Supplementary File 2 – Scaffolds fasta sequences in which homeodomains were detected (as described in Table S4).

## Notes

https://doi.org/10.6084/m9.figshare.13333841

### Competing Interest Statement

The authors have declared no competing interest.

